# Testing for association with rare variants in the coding and non-coding genome: RAVA-FIRST, a new approach based on CADD deleteriousness score

**DOI:** 10.1101/2021.11.04.467235

**Authors:** Ozvan Bocher, Thomas E. Ludwig, Gaëlle Marenne, Jean-François Deleuze, Suryakant Suryakant, Jacob Odeberg, Pierre-Emmanuel Morange, David-Alexandre Trégouët, Hervé Perdry, Emmanuelle Génin

## Abstract

Rare variant association tests (RVAT) have been developed to study the contribution of rare variants widely accessible through high-throughput sequencing technologies. RVAT require to aggregate rare variants in testing units and to filter variants to retain only the most likely causal ones. In the exome, genes are natural testing units and variants are usually filtered based on their functional consequences. However, when dealing with whole-genome sequence (WGS) data, both steps are challenging. No natural biological unit is available for aggregating rare variants. Sliding windows procedures have been proposed to circumvent this difficulty, however they are blind to biological information and result in a large number of tests.

We propose a new strategy to perform RVAT on WGS data: “RAVA-FIRST” (RAre Variant Association using Functionally-InfoRmed STeps) comprising three steps. (1) New testing units are defined genome-wide based on functionally-adjusted Combined Annotation Dependent Depletion (CADD) scores of variants observed in the GnomAD populations, which are referred to as “CADD regions”. (2) A region-dependent filtering of rare variants is applied in each CADD region. (3) A functionally-informed burden test is performed with sub-scores computed for each genomic category within each CADD region. Both on simulations and real data, RAVA-FIRST was found to outperform other WGS-based RVAT. Applied to a WGS dataset of venous thromboembolism patients, we identified an intergenic region on chromosome 18 that is enriched for rare variants in early-onset patients and that was that was missed by standard sliding windows procedures.

RAVA-FIRST enables new investigations of rare non-coding variants in complex diseases, facilitated by its implementation in the R package Ravages.

**Author Summary:** Technological progresses have made possible whole genome sequencing at an unprecedented scale, opening up the possibility to explore the role of genetic variants of low frequency in common diseases. The challenge is now methodological and requires the development of novel methods and strategies to analyse sequencing data that are not limited to assessing the role of coding variants. With RAVA-FIRST, we propose a novel strategy to investigate the role of rare variants in the whole-genome that takes benefit from biological information. Especially, RAVA-FIRST relies on testing units that go beyond genes to gather rare variants in the association tests. In this work, we show that this new strategy presents several advantages compared to existing methods. RAVA-FIRST offers an easy and straightforward analysis of genome-wide rare variants, especially the intergenic ones which are frequently left behind, making it a promising tool to get a better understanding of the biology of complex diseases.

## Introduction

With advance in sequencing technologies, it is now possible to explore the role of rare genetic variants in complex diseases. Different rare variant association tests (RVAT) have been developed that gather rare variants into testing units and compare rare variant content in these testing units between cases and controls (1–3). While the impact of rare variants has already been shown in several complex diseases (4–6), RVAT face two key challenges: (i) the definition of the testing units and (ii) the selection of the qualifying rare variants to include in these units. The proportion of causal variants in the testing units being a major driver of power, especially for burden tests, it is indeed important to ensure that qualifying variants are enriched in variants likely to have some functional impact (3,7). When exome analyses are undertaken, rare variants are most often grouped by genes and included in the analysis depending on their impact on the corresponding protein (8,9). Nevertheless, the gene definition is not always optimal as differences in rare variants burden between cases and controls could sometimes only be found in a sub-region of a gene. This is for example the case in the *RNF213* gene where an enrichment in rare variants located in the C-terminal region is found in Moyamoya cases (10). Defining testing units and qualifying variants is also much more challenging in the non-coding genome due to the lack of defined genomic elements and the higher difficulty to predict the functional impact of non-coding variants (11). It is yet a question of interest as several studies have shown the importance of rare non-coding variants in the development of complex diseases (12–14). Functional elements such as enhancers or promoters can be used as testing units (5,15,16) but they prevent the analysis of all rare variants in the genome and can be too small to get a sufficient number of rare variants for association analysis. On the other hand, sliding windows procedures such as SCAN-G (17) or WGSCAN (18) can be used to test for association over the whole genome. Nevertheless, they present several limits including the window definition that is arbitrary and blind to biological information, the high number of tests and the associated computation time. With overlapping windows, there is also a strong correlation between tests performed in the different testing units that requires the use of permutation procedures to account for multiple testing. Finally, to filter rare variants in the testing units, pathogenicity scores are often used but without guidelines on which score to use and which threshold to apply.

In this paper, we propose RAVA-FIRST (RAre Variant Association using Functionally InfoRmed STeps), a new strategy for analysing rare variants in the coding and the non-coding genome that addresses the previous issues. First, we provide pre-defined testing units in the whole genome called “CADD regions” based on the Combined Annotation Dependent Depletion (CADD) scores of deleteriousness of variants observed in the GnomAD general population. These regions prevent the use of sliding windows procedures while enabling the study of rare variants in the whole genome. Second, we propose a filtering approach based on CADD scores with region-dependant thresholds to represent the genetic context of each CADD region and avoid the use of a fix threshold along the genome. Finally, we integrate functional information into the burden test to detect an accumulation of rare variants in specific genomic categories within CADD regions. Through a statistical description of these testing units, we show that they preserve the integrity of the majority of functional elements in the genome. We also show that the RAVA-FIRST filtering strategy enables a better discrimination between functional and non-functional variants within the testing units. We applied RAVA-FIRST to real whole-genome sequencing data from individuals with venous thromboembolism (VTE) and detected an intergenic association signal that would have been missed with sliding windows and a classical filtering of rare variants. RAVA-FIRST is implemented in the R package Ravages available on the CRAN and maintained on github (19,20).

### Description of the Method

RAVA-FIRST is developed to test for association with rare variants in the whole genome. It deals with all steps from the definition of testing units and the filtering of rare variants, to the association test accounting for functional information. The main steps are represented in S1 Fig and further details are presented hereafter.

### Testing units in RAVA-FIRST: the CADD regions

Following Havrilla et al. (2019) (21), we seek to identify some genomic regions that were significantly depleted in functional variants to use them as testing units in RVAT. For that purpose, Havrilla et al. (2019) defined “constrained coding regions” (CCR) as exonic regions where no important functional variation (defined as being at least missense) was found in the general population of GnomAD (22). In our experience, two limits prevent the direct use of CCR as testing units in the whole genome: they are too small to gather a sufficient number of rare variants (224 bp being the maximum length of a CCR) and their definition relies on the consequence of the variants on the translated protein, not available in the non-coding genome. We therefore decided to expand the proposed approach by estimating the functionality of variants through CADD scores (23). CADD scores were chosen because of their availability for every substitution in the genome and because they rank well in the comparison test of functional annotation tools (24).

Coding variants tend to present higher CADD values than non-coding variants (23). A selection based on a CADD threshold would therefore result in a majority of coding variants selected. In order to avoid this pattern, we adjusted the RAW CADD scores on a PHRED scale within each of three genomic categories: “coding”, “regulatory” and “intergenic” regions. Coding regions correspond to CCDS (25) and represent 1.2% of the genome. Regulatory regions represent 44.3% of the genome and are defined by the union of introns, 5’ and 3’ UTR, promoters and enhancers, all being involved in gene regulation (26). Enhancers and promoters have been obtained with the SCREEN tool from ENCODE which enables the definition of a large number of regulatory elements in diverse cell types (27). Finally, intergenic regions correspond to all regions not being described as coding or regulatory regions, representing 54.5% of the genome. More details are given in the Supporting Information.

Adjusted CADD scores were used to select the variants that will bound the “CADD regions”. First, we selected the variants with an adjusted CADD score greater than 20, that is the top 1% of variants with the highest predicted functional impact within each of the three genomic categories. Then, among those variants, only the ones observed at least two times in GnomAD r2.0.1 genomes were further selected and used as boundaries of CADD regions. For CADD regions to be used as testing units in RVAT, they need to be large enough to contain several rare variants. Contiguous small regions of less than 10 kb were therefore grouped together to form clusters of variants with high adjusted CADD scores. Non-sequenced regions and low-covered regions in GnomAD containing potential important functional variants were excluded from CADD regions, leading to gaps within CADD regions of at least one base pair (i.e. no CADD region overlap them to avoid artificially long regions due to a lack of variants in GnomAD). Finally, CADD regions are only defined for regions where CADD scores are available (removing among others centromeres and telomeres). Note that CADD regions can overlap different genomic categories (coding, regulatory or intergenic). More details about the steps and parameters used for the definition of CADD regions are presented in the Supporting Information.

### The RAVA-FIRST filtering strategy

In addition to the definition of new testing units in the whole genome, we propose a new filtering strategy in RAVA-FIRST to select qualifying variants. Using gene-specific CADD thresholds rather than a fixed threshold for all genes was previously found to improve prediction (28). Building on the same idea, we defined thresholds that are specific to each CADD region. To define these region-specific thresholds, we derived the median of all adjusted CADD scores of variants observed at least two times in GnomAD in each CADD region. This value represents the median score level that is tolerated in the general population within each CADD region. Adjusted CADD scores refer here to the PHRED CADD scores computed respectively for coding, regulatory and intergenic genomic categories as defined before. Qualifying variants are then defined as rare variants with an adjusted CADD score above the threshold specific to their region. Note that because CADD scores are only available for SNVs, other types of variants are excluded from the analyses.

### Burden test in RAVA-FIRST: taking into account functional information

As mentioned before, several of the CADD regions overlap different genomic categories (coding, regulatory or intergenic, Figs S1 and S3). As the effects of variants belonging to these different genomic categories may not be the same, we extended the burden test defined as:

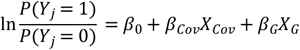

With *Y*_*j*_ the vector of phenotypes for the n individuals: 0 for the group of controls and 1 for the group of cases. *β*_0_ represents the intercept of the model and *X*_*Cov*_ the matrix of covariates (if any) with their associated effect, *β*_*Cov*_. *β*_*G*_ corresponds to the estimated effect of the burden *X*_*G*_, computed for example using WSS (1) which corresponds to a weighted sum of rare alleles based on their frequency, the rarest alleles having the highest weights.

To take into account functional information, we integrated a sub-score for each genomic category into the regression model, similarly to the analysis of rare and frequent variants proposed by Li and Leal (2008) (7):

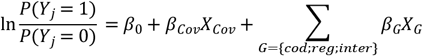

Sub-scores *X*_*G*_ are constructed for each genomic category within a CADD region, with at most three sub-scores (coding, regulatory or intergenic). The p-value can be determined using a likelihood ratio test comparing this model to the null model where the sub-scores are not included. This sub-score analysis, also called RAVA-FIRST burden test, is also available for continuous and for categorical phenotypes using the extension of burden tests developed in Bocher et al. (2019) (19). The RAVA-FIRST burden test coupled with the region-specific filtering on the adjusted CADD score enables to keep the most important functional variants within each genomic category and to take into account those categories in the association test while performing only one test by CADD region.

### Verification and Comparison

#### Statistics on CADD regions and comparison with genomic elements

A total of 135,224 CADD regions were defined covering 93.2% of the genome (in build GRCh37). Among CADD regions, several are very small in size, despite our approach to combine small regions, due to the removal of low-covered regions, preventing their use in RVAT. We therefore decided to focus on the 106,251 CADD regions larger than 1kb, which cover 93% of the genome. Among those CADD regions, 28.3% span only one type of genomic category, 58.5% span two of the three types of genomic categories, and 13.2% overlap the three genomic categories (S3 Fig). Some CADD regions are extremely large, mainly around the centromeres (Table 1). About 80% of CADD regions have a size between 5 and 50 kb with a mean of 25 kb, making them completely compatible with the size of genes commonly used as testing units used in RVAT.

**Table 1:** Summary statistics of the lengths of CADD regions (larger than 1 kb)

|  | Quantiles |  |  |  |  | Mean |
| --- | --- | --- | --- | --- | --- | --- |
|  | 0% | 25% | 50% | 75% | 100% |  |
| Length (kb) | 1 | 10.790 | 16.579 | 29.116 | 1,731.228 | 25.224 |

We then compared the position of genomic elements relative to the defined CADD regions (Table in S1 Table shows how the different genomic elements have been obtained). A large majority of genomic elements are entirely included into a single CADD region and thus their integrity is preserved (Table 2). This is expected as all these genomic elements are substantially smaller than the CADD regions and therefore have a high probability of being included in a CADD region. For larger elements such as introns or lncRNA, the percentage decreases but remains high (more than 80% of lncRNA are overlapped by at most 2 CADD regions). The genomic elements spanning more than one CADD region are on average longer than the ones being entirely included into a single CADD region. However, when comparing CCR and CADD regions, it is interesting to note that the CCRs entirely encompassed within a single CADD region are the longest ones that should also represent the most constrained regions.

**Table 2:** Percentage of genomic elements entirely encompassed within a CADD region

| Exon<br>CCDS | Protein<br>domains | CCR | Introns/UTR | Enh-Prom |  | Silencers | CTCF | lncRNA |
| --- | --- | --- | --- | --- | --- | --- | --- | --- |
|  |  |  |  | DECRES | ENCODE |  |  |  |
| 97.8% | 81.8% | 99.2% | 85.9% | 93.1% | 96.4% | 95.1% | 95.8% | 65.5% |

#### Performance of RAVA-FIRST filtering based on adjusted CADD scores

To assess the performance of the adjusted CADD scores and the RAVA-FIRST filtering, we evaluated its capacity in discriminating benign from pathogenic variants using the Clinvar database (29). We computed true positive rate (TPR), true negative rate (TNR) and precision for the RAVA-FIRST filtering and compared the results to the ones obtained by applying a fixed CADD threshold of 10, 15 or 20 on variants annotated with CADD scores v1.4. After the selection of rare variants included in RVAT (see the Supporting Information), the dataset of analysis contains 70,931 variants of which 25,931 are benign and 45,000 are pathogenic. All filtering strategies show a very high TPR (Fig 1A), meaning that the majority of pathogenic variants would be selected as qualifying variants for RVAT. The TNR increases with the increasing CADD score threshold which is expected as less variants, and therefore less benign variants, are included in the analysis. The RAVA-FIRST filtering shows the highest TNR and the highest precision. While the TPR value is extremely important to select the most probable causal variants in RVAT, it is also important to have a high TNR value, otherwise the signal will be diluted by a high proportion of non-causal variants. The precision value summarises the TPR and TNR parameters and therefore, to a certain extent, is representative of the percentage of causal variants among selected variants. Therefore, we show that the RAVA-FIRST filtering strategy is the most accurate to select qualifying rare variants for RVAT. Focusing on the coding genome, we also compared the performance of RAVA-FIRST filtering approach against two others approaches classically used on genes as testing units: (1) filter for variants with a functional impact expected to change the protein (“missense_variant”, “missense_variant&splice_region_variant”, “splice_acceptor_variant”, “splice_donor_variant”, “start_lost”, “start_lost&splice_region_variant”, “stop_gained”, “stop_gained&splice_region_variant”, “stop_lost”, “stop_lost&splice_region_variant” and “stop_retained_variant”), and (2) filter on the MSC value, a gene-specific CADD threshold(28). These two filtering approaches resulted in a slightly higher TPR than our proposed strategy but lower TNR and lower precision (Fig 1B). Therefore, even in an exome analysis, the RAVA-FIRST filtering outperforms classical filtering strategies to select qualifying rare variants for RVAT.

**Figure 1:**
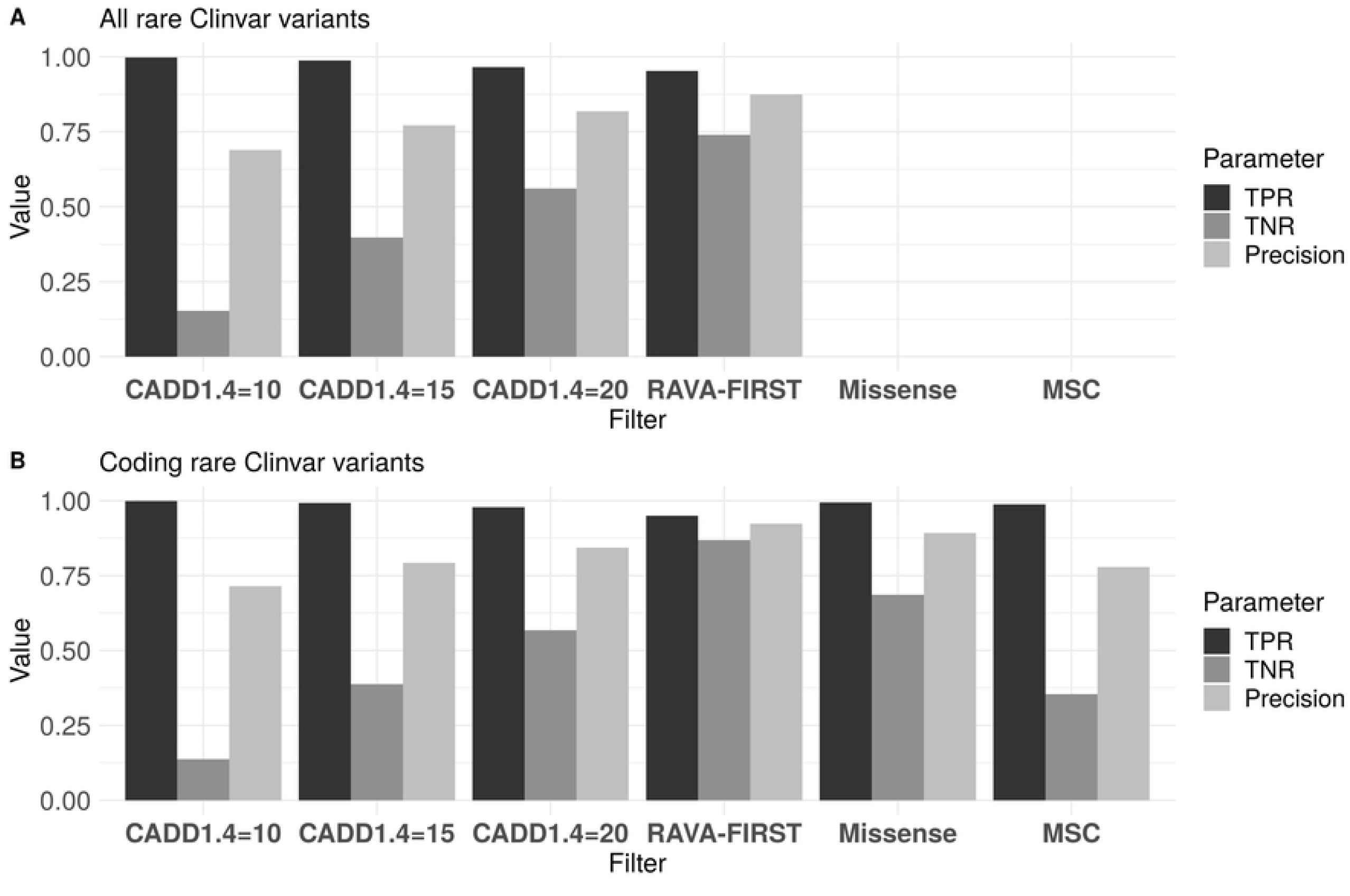
TPR, TNR and precision of different filtering strategies on the whole Clinvar dataset or only Clinvar coding variants.

Finally, we investigated the performances of these different strategies on different classes of non-coding variants (S4 Fig). All the performances are lower than in the coding genome, especially the TPR that is much lower for strategies based on a fixed CADD threshold, highlighting the fact that CADD values are lower in the non-coding genome and adjusted CADD threshold may therefore be preferred. RAVA-FIRST filtering using region-dependant thresholds keeps the highest precision in the different classes of variants, except for UTR variants where a slight decrease of TNR and precision is observed. Note however that these results may not be as accurate as those obtained on the coding regions as much fewer variants are included: 2,309, 4,048 and 617 for UTR, introns and intergenic variants respectively compared to 54,664 coding variants.

### RAVA-FIRST burden test – Simulations

To validate the RAVA-FIRST burden test, we performed simulations under the null hypothesis and under different scenarios of association using data from the 1000 Genomes European populations (30) in the *LCT* gene. We simulated 1,000 controls and 1,000 cases using the simulations based on haplotypes implemented in the R package Ravages (19). A total of 201 variants was considered in the *LCT* gene. These variants were polymorphic in the European populations and rare variants were defined with a MAF lower than 1%. Two CADD regions overlap the *LCT* gene, R019233 and R019234, containing respectively 75 and 126 variants, both overlapping coding and regulatory categories.

#### Type I error

We first simulated data under the null hypothesis to verify that the RAVA-FIRST burden test maintains appropriate type I errors. We simulated two groups of 1,000 individuals in the R019234 CADD region without any genetic effect and we applied the classical WSS and the RAVA-FIRST WSS. Type I errors were computed using 5·10^6^ simulations at three significance levels: 5·10^−2^, 10^−3^ and 2.5·10^−6^ (the usual threshold for whole exome rare variant association tests). The RAVA-FIRST WSS maintains good type I error levels at these different significance thresholds, similar to the ones obtained with the classical WSS (Table in S2 Table).

#### Power analysis

We then performed a power study based on simulations at two levels: at the level of the R019234 CADD region and at the level of the *LCT* gene. In both cases, we simulated 50% of causal variants randomly in the whole unit (scenarios S1 and S3), in the coding regions (scenarios S2A and S4A) or in the regulatory regions (scenarios S2B and S4B). All the scenarios are summarised in Table 3. We compared the classical WSS to the RAVA-FIRST WSS using the gene or the two CADD regions as testing units. When CADD regions were used as testing units, analyses were performed for each of the two CADD regions and the minimum p-value was taken and multiplied by two to correct for multiple testing. A total of 1,000 replicates were simulated for each scenario and power was assessed at a genome-wide significance threshold of 2.5·10^−6^.

**Table 3:** Scenarios of association simulated to assess the performance of the RAVA-FIRST burden test

|  | *LCT* gene |  |  |  |
| --- | --- | --- | --- | --- |
|  | R019233 |  | R019234 |  |
|  | Coding | Regulatory | Coding | Regulatory |
| S1 |  |  | 50% |  |
| S2A |  |  | 50% | 0% |
| S2B |  |  | 0% | 50% |
| S3 | 50% |  |  |  |
| S4A | 50% | 0% | 50% | 0% |
| S4B | 0% | 50% | 0% | 50% |

Table 4 presents the power results obtained from this simulation study for both the classical WSS and the RAVA-FIRST WSS. Similar trends were observed between the two analyses, regardless if the simulations are performed at the scale of CADD regions or at the scale of the gene. When the causal variants were randomly sampled across the entire region (scenarios S1 and S3), the classical WSS with only one score for the entire region slightly outperformed the RAVA-FIRST method with sub-scores. Nevertheless, the loss of power for the latter was modest (less than 10%). By contrast, when causal variants were present only in the coding categories (scenarios S2A and S4A), which represent a small proportion of the entire region (approximately 15%), the RAVA-FIRST strategy was much more powerful than the classical WSS (approximately 50% gain in power). When causal variants were present in the regulatory categories only (scenarios S2B and S4B), both strategies showed similar power. All these results highlight the gain of power using the RAVA-FIRST WSS when a cluster of causal variants is present within a functional category of the CADD region while maintaining good power levels when causal variants are spread all across the region. When comparing the simulations with causal variants sampled at the gene level or at the CADD region level, burden tests gathering variants within the corresponding testing units show, as expected, the highest levels of power. Nevertheless, the loss of power when using CADD regions as testing units instead of the entire gene is lower when causal variants are sampled across the entire gene (scenario S3) than the gain of power they present when causal variants are sampled within a specific CADD region (scenario S1). This is particularly true for the RAVA-FIRST WSS.

**Table 4:** Power at the genome-wide significance level of 2.5·10^−6^ under the different simulation scenarios using either the classical WSS or the RAVA-FIRST WSS at the scale of either the entire gene or CADD regions

|  | By gene |  | By CADD regions |  |
| --- | --- | --- | --- | --- |
|  | Classical WSS | RAVA-FIRST WSS | Classical WSS | RAVA-FIRST WSS |
| S1 | 0.409 | 0.370 | 0.782 | 0.701 |
| S2A | 0 | 0.431 | 0.002 | 0.602 |
| S2B | 0.408 | 0.404 | 0.689 | 0.706 |
| S3 | 0.751 | 0.678 | 0.512 | 0.433 |
| S4A | 0.004 | 0.564 | 0.012 | 0.474 |
| S4B | 0.657 | 0.64 | 0.39 | 0.391 |

### Applications

#### Ethics Statement

The MARTHA study was approved by its institutional ethics committee and informed written consent was obtained in accordance with the Declaration of Helsinki. Ethics approval were obtained from the “Departement santé de la direction générale de la recherche et de l’innovation du ministère” (Projects DC: 2008-880 and 09.576).

### RAVA-FIRST analysis

RAVA-FIRST was used on whole genome sequence (WGS) data from patients affected by venous thromboembolism (VTE). VTE is a multifactorial disease with a strong genetic component (31). There exists a huge heterogeneity between patients in the age at first VTE event. To study the role of rare variants on VTE age of onset, WGS data were used from 200 individuals from the MARTHA cohort (32). These individuals were selected among patients with unprovoked VTE event who were previously genotyped for a genome-wide association study (33) and present no known genetic predisposing factor. Individuals were dichotomized based on the age at first VTE event either before 50 years of age (early-onset) or after (late-onset). The threshold of 50 years was chosen based on the results of recent studies (34) that hint toward a genetic heterogeneity between these two groups. A quality control (QC) of the sequencing data was performed using the program RAVAQ (https://gitlab.com/gmarenne/ravaq). After QC, 184 individuals were included for analysis with 127 presenting an early-onset VTE and 57 a late-onset VTE. Only variants passing all QC steps and with a MAF lower than 1% in the sample were considered in the association tests comparing early and late-onset groups. For these comparisons, rare variants were gathered either by CADD regions or by using the sliding windows procedure implemented in WGScan (18). Qualifying variants were selected based on CADD scores and using two filtering strategies: a fixed CADD threshold of 15 (as recommended by https://cadd.gs.washington.edu/info, version v1.4) or the RAVA-FIRST CADD region-specific filtering (applied on adjusted scores). Association was tested using the WSS burden test. When the RAVA-FIRST filtering was used, the corresponding WSS test with sub-scores was applied. Table 5 shows the number of testing units and variants kept under each strategy. For all tests with CADD regions, only regions containing at least 5 rare variants were kept. WGScan was used with default parameters, i.e. with testing units of 5, 10, 15, 25 or 50 kb.

**Table 5:** Number of testing units and variants kept under the three strategies

|  | Testing units | Filtering | Number of testing units | Number of variants |
| --- | --- | --- | --- | --- |
| WGSscan<br>Fixed CADD threshold | Sliding windows | $MAF \leq 1\%$<br>$CADD \ v1.4 \geq 15$ | 377,092 | 96,347 |
| RAVA-FIRST units<br>(CADD regions)<br>Fixed CADD threshold | CADD regions | $MAF \leq 1\%$<br>$CADD \ v1.4 \geq 15$ | 10,389 | 96,294 |
| RAVA-FIRST units<br>(CADD regions)<br>RAVA-FIRST filtering | | $MAF \leq 1\%$<br>Adjusted CADD ≥ median | 95,690 | 3,641,502 |

QQ-plots for the WSS tests using those three strategies are shown in Fig 2. As expected, a lower significance threshold is required to reach genome-wide significance with the sliding window procedure due to the higher number of testing units. Accordingly, the computation time was much lower for the two analyses by CADD regions (6min when filtering based on a fixed CADD score threshold and 25min when using the region-specific CADD thresholds) than for the sliding windows procedure (47min). Our dataset contains less than 200 individuals, suggesting that the gain in computation time of CADD regions compared to sliding window procedures would be even greater in larger WGS datasets. No significant result was found when selecting variants with a CADD score greater than 15 using neither the sliding window strategies nor the CADD regions to gather rare variants, whereas one association reached borderline significance (p = 6.41·10^−7^) when using the RAVA-FIRST strategy.

**Figure 2:**
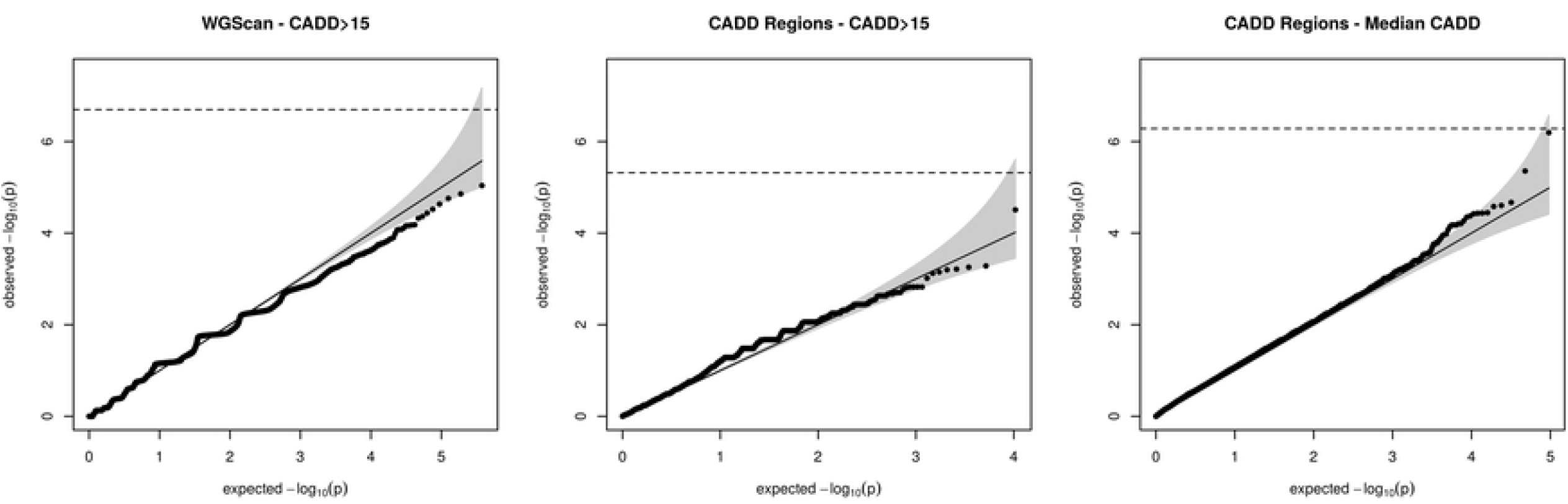
QQ-plot of WSS analyses on VTE data using the three strategies of analysis. Early-onset patients (<50 years old) were compared to late-onset patients (≥50 years old).

This association maps to R126442, a CADD region of 21 kb on chromosome 18:66788277-66809402 that contains 31 rare variants after RAVA-FIRST filtering. In this region, none of the variants observed in VTE patients or in GnomAD achieved a CADD score above 15. This explains why the association could not have been detected by the two other strategies based on fixed CADD score ≥ 15. The median of CADD scores observed for GnomAD variants in this region is 1.44 and the adjusted CADD scores of selected variants range from 1.62 to 8.50. These observations emphasize the need to adapt thresholds depending on the genomic region under analysis. Interestingly, only early-onset VTE patients carry qualifying rare variants and have non-null WSS scores (Fig 3). Among early-onset patients, a trend is also observed for WSS scores to decrease with increasing age of onset.

**Figure 3:**
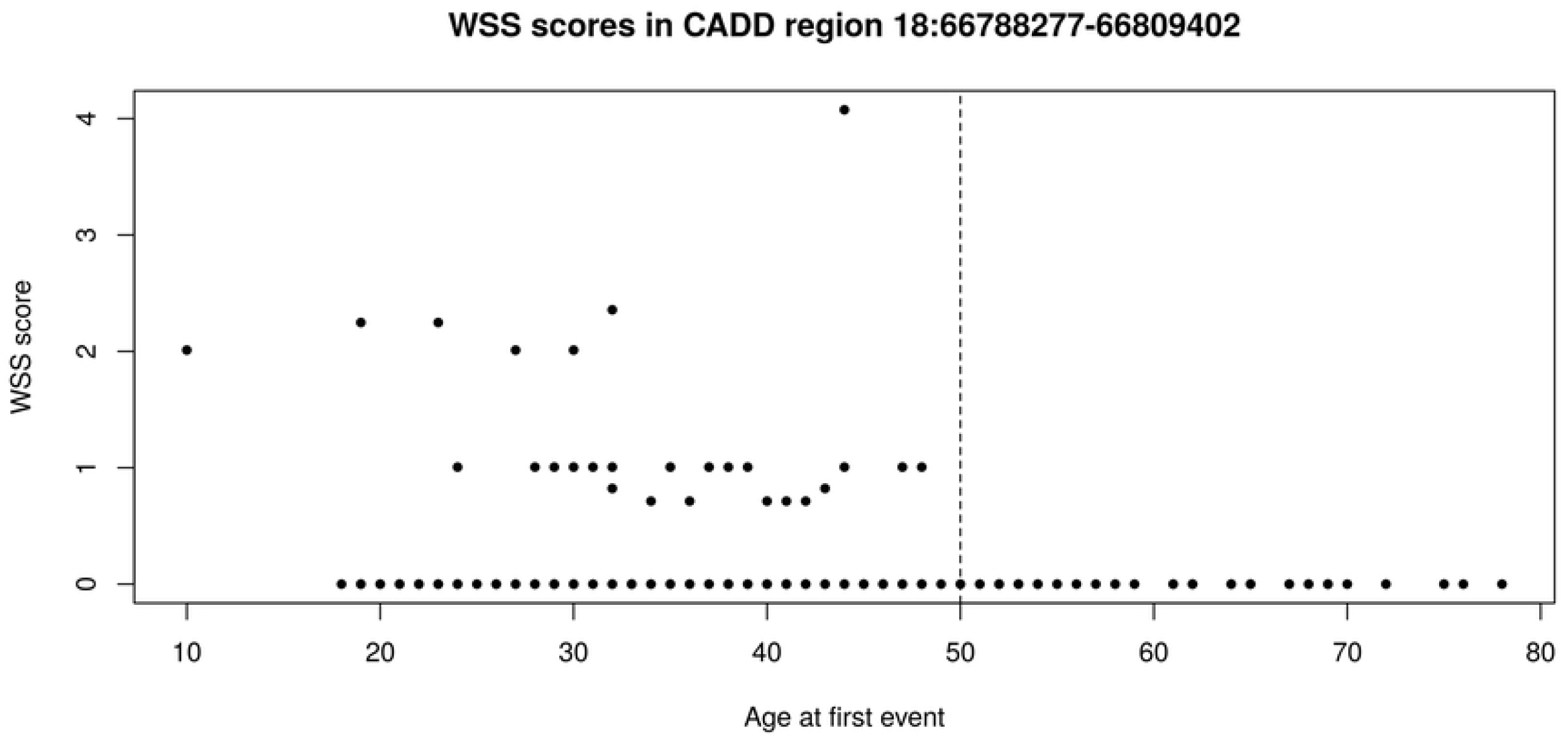
WSS scores in the CADD region depending on the age at first VTE event. The dashed line corresponds to the age 50 discriminating early onset from late onset events.

The CADD region R126442 was then tested for association with 20 biological VTE biomarkers available in MARTHA patients: antithrombin, basophil, eosinophil, Factor VIII, Factor XI, fibrinogen, hematocrit, lymphocytes, mean corpuscular volume, mean platelet volume, monocytes, neutrophils, PAI-1, platelets count, protein C, protein S, prothrombin time, red blood cells count, von Willebrand Factor, and white blood cells count. For this, a linear regression model was used where adjustment was made on age at sampling and sex. At the Bonferonni threshold of 0.0025, one significant association (p = 7.1·10^−4^) was observed, VTE patients with a non-null WSS score exhibiting decreased haematocrit levels, a surrogate marker of red blood cells (Table in S3 Table). A similar trend (p = 4.6·10^−3^) was observed with red blood cell count.

We also investigated the association of the identified region with 376 plasma protein antibodies that were selected to be involved in thrombosis-related processes and that have been previously profiled in MARTHA (32,35). Regression analysis were conducted on log transformed values of antibodies and were adjusted for age, sex, and three internal control antibodies. In order to handle the correlation between measured protein antibodies, we used the Li and Ji method (36) to estimate the number of effective independent tests. This number, calculated to be 163, was then used to define a Bonferroni threshold for declaring study-wide statistical significance. While not reaching the study-wise significance level of p = 3.1·10^−4^ after correction for multiple testing, it is worth noting that the two proteins that exhibited the strongest significance with marginal association at p < 0.001, procalcitonin tagged by the HPA043700 antibody (p = 7.2·10^−4^) and PDPK1 tagged by HPA035199 (p = 7.5·10^−4^), have been both proposed to be involved in red blood cell biology (37,38).

According to ENCODE data, the R1246442 CADD region overlaps “intergenic” and “regulatory” categories with one distant enhancer-like signature. To describe this region further, we looked at TADs positions in https://dna.cs.miami.edu/TADKB/brows.php in HUVEC and HMEC cell lines, two cell types known to be relevant for VTE pathophysiology. We found that the CADD region is included into the topological associated domains (TADs) 18:66450000-68150000. By studying TADs described by Lieberman-Aiden et al. 2009 in other cell lines such as KBM7, K562 or GM12878, we retrieved a TAD with similar positions, giving additional evidence for the presence of this TAD around the CADD region associated with early-onset patients. We then explored this TAD region for the presence of candidate VTE genes whose regulation could be influenced by the enhancer region that maps our R1246442 region. Using the UCSC genome browser (40) integrating information about interactions between GeneHancer regulatory elements and genes expression (see S5 Fig), we identified *CD226* as a strong biological candidate. *CD226* codes for a glycoprotein expressed at the surface of several types of cells, including blood cell, and several studies have shown that it was associated with vascular endothelial dysfunction (41–43). Genetic variants in *CD226* have also been found associated with several blood cell traits including platelets, white blood cells (e.g. neutrophil, eosinophil) (44) and reticulocyte counts (45), another red blood cell biomarker.

## Discussion

Even though whole genome sequencing data are now more often available on cases and controls, rare variant association tests (RVAT) usually remain restricted to the coding part of the genome. This is explained by the lack of tools to explore rare variant associations outside genes (11). Indeed, RVAT requires the definition of testing units that are easily defined through genes in coding regions and the selection in these regions of the most functionally-relevant variants. This is also easier in the coding genome as most prediction tools were developed and tested through the effects of variants on encoded proteins. In the non-coding genome, testing units can be defined based on functional elements such as enhancers or silencers, or through the use of sliding window procedures. The first solution prevents RVAT from being applied to all rare variants in the genome as biological units are not defined over the entire genome. The second strategy with sliding windows results in a large number of tests and the need to adjust p-values to take into account the multiple correlated tests performed. In this work, we propose an entire new strategy of analysis of rare variants in the coding and the non-coding genome, RAVA-FIRST, which is composed of three steps. Firstly, RAVA-FIRST proposes some new testing units to gather rare variants, the so-called “CADD regions” that we defined over the entire genome based on CADD scores of variants observed in GnomAD. These CADD regions are large enough to include a sufficient number of rare variants to allow RVAT. They tend to preserve functional elements that, for a majority of them, are not split into several CADD regions. Secondly, RAVA-FIRST filters variants based on region-specific adjusted CADD thresholds that allow to select the best candidate variants within each region. This filtering approach was found to be more efficient than traditional approaches to discriminate between benign and pathogenic variants within a set of variants. Indeed, our benchmarking study using a set of Clinvar variants showed that the other filtering strategies we considered were good at identifying true causal variants (true positive rates were high) but bad at finding the non-causal variants (true negative rates were low). Both true positive and true negative rates are important to achieve a high percentage of causal variants within testing units, this percentage being the main driver of power in RVAT, especially in burden tests (2,3,7). Thus, the RAVA-FIRST filtering strategy is expected to result in an appreciable increase of power as compared to classically used strategies. Indeed, RAVA-FIRST enables to keep the most important functional variants within coding, regulatory and intergenic categories of the genome by adapting CADD score threshold to the genomic context. Finally, RAVA-FIRST includes a burden test that integrates information on genomic categories in the regression and that, coupled with the region-specific filtering, leads to a better detection of causal variants, should they cluster in one of these genomic categories only. We also showed through simulations that good power levels were maintained using RAVA-FIRST burden test when causal variants were randomly sampled.

RAVA-FIRST was applied on real WGS data from VTE patients where an accumulation of rare variants in patients with early-onset events was investigated. We did not detect any significant signal using the sliding window procedure or CADD regions when qualifying rare variants were selected based on a fixed CADD threshold. However, we detected an association signal using both the grouping and filtering of rare variants proposed in RAVA-FIRST. The associated CADD region is intergenic, contains a predicted enhancer and is surrounded by a TAD containing 5 genes including *CD226*, a strong candidate for blood cell traits that are new well recognized to be key players in VTE physiopathology (31). All rare variants in this region present low CADD scores and were not even included in analyses based on a fix CADD threshold, highlighting the importance of taking into account the genetic context to detect the most important predicted functional variants within each CADD region. These 31 rare variants are exclusively observed in early-onset cases. Fourteen of these variants are absent from GnomAD, and 10 of the 17 remaining variants have a lower frequency in GnomAD population than in our sample. This reinforces the value of the association signal in this CADD region, although it should be further described and validated using functional experiments. Preliminary investigations that need to be further explored, at both experimental and epidemiological levels, strongly suggest that this region is associated with several inflammatory markers impaired in anaemia of inflammation (38,46) and in platelets, both mechanisms being involved in thrombotic processes (47).

Some limits can be pointed out on our RAVA-FIRST approach. Firstly, the definition of CADD regions relies on the GnomAD population and on the adjusted CADD threshold. We chose to use the whole GnomAD dataset but it could be of interest to select some of the populations to be more specific. It has for example been suggested that different expression patterns could be found between different populations (48). Nevertheless, in classical exome analyses, rare variants are mostly filtered based on the maximum frequency observed among multiple populations. Furthermore, CADD regions are not defined for low-covered and non-sequenced genomic regions in GnomAD and their definition could therefore be improved in the future. Concerning the definition of the genomic categories, we decided to include all genomic elements directly implicated into regulatory functions to define the regulatory regions of the genome, but we did not include silencers or lncRNA for example. However, the choice of elements to include as the regulatory category will only impact the adjusted CADD scores that are similar between regulatory and intergenic regions, and won’t therefore have a huge impact on CADD regions definition. As an example, using DECRES (49) to predict enhancers and promoters instead of SCREEN results in a very high correlation between the definition of CADD regions, 80% of them being identical.

On the other hand, the pre-definition of regions in the whole genome offers several advantages, including the region-specific filtering mentioned before. In addition, the newly defined CADD regions can be used in existing software that require regions as input parameters (50,51), enabling to apply a wide variety of RVAT available in those programs to the whole genome. Especially, Bayesian methods which have been shown to be of great promise in the analysis and filtering of rare variants (52,53) could be applied beyond genes by using CADD regions.

To our knowledge, CADD regions represent predefined testing units for RVAT that cover the highest proportion of the genome. These regions have been made publicly available (cf “Data availability” section below). CADD regions are part of a whole new strategy of rare variant analysis in the whole genome, RAVA-FIRST, that further benefits from the integration of functional information both for the filtering of rare variants and their analysis with burden tests. RAVA-FIRST has been implemented in the package R Ravages available in the CRAN and on Github, offering an easy and straightforward tool to perform RVAT in the whole genome. We believe that our developments will help researchers to explore the role of genome-wide rare variants in complex diseases. Firstly, through the redefinition of testing units in the coding genome where cluster of causal variants can be found within genes and retrieved using CADD regions (10). Secondly, through the study of non-coding variants, especially intergenic ones, which are currently often excluded from the analysis. Going beyond the gene and the consequences on proteins, RAVA-FIRST will help for a better understanding of biological mechanisms behind complex diseases.

## Data availability

The files containing the positions of CADD regions, the positions of genomic categories and the adjusted CADD scores are available at https://lysine.univ-brest.fr/RAVA-FIRST/. All the functions needed for RAVA-FIRST to annotate, group, filter and analyse rare variants have been implemented in the package R Ravages (https://cran.r-project.org/web/packages/Ravages/, https://github.com/genostats/Ravages) which directly downloads the files from https://lysine.univ-brest.fr/RAVA-FIRST/.

Information about the CADD region R126442 that was found associated with VTE age at first event is available in the Supporting Information File 2. Information about individuals (WSS score, age and sex) and variants (position, adjusted CADD score and weight in WSS) are given.

## Supporting Information captions

**S1 Fig. Steps performed in RAVA-FIRST**.

**S2 Fig. Definition of CADD regions and removal of low-covered and non-sequenced regions in GnomAD**.

**S3 Fig. Percentage of CADD regions (≥1kbp) overlapping each of the three genomic categories**.

**S4 Fig. TPR, TNR and precision of different filtering strategies on Clinvar non-coding variants (UTR, introns or intergenic regions)**.

**S5 Fig. Screenshot of the TAD 18:66450000-68150000 in the UCSC genome browser containing the CADD region R126442 and a potential enhancer regulating the CD226 gene, a candidate gene in VTE**.

**S1 Table. Sources used to get genomic elements for comparisons with CADD regions**.

**S2 Table. Type I error of the classical WSS and the RAVA-FIRST WSS using 5·10**^**6**^ **simulations under the null hypothesis**.

**S3 Table. Characteristics of the studied VTE sample**. Mean (Standard Deviation) for quantitative variables. Count (%) for qualitative variables.

**S1 File. Details about the RAVA-RIST method and its evaluation**.

**S2 File. Information on the CADD region R126442 associated with VTE age at onset**. Information about individuals (WSS score, age and sex) and variants (position, adjusted CADD score and weight in WSS) are given.

